# *Coxiella burnetii* establishes a small cell variant (SCV)-like persistent form to survive adverse intracellular conditions

**DOI:** 10.1101/2025.09.18.677042

**Authors:** Faiza Asghar, Inaya Hayek, Christian Berens, Elisabeth Liebler-Tenorio, Anja Lührmann

## Abstract

*Coxiella burnetii* is an obligate intracellular zoonotic bacterium that causes Q fever. Infections can be either acute or chronic. Of note, chronic Q fever develops months or years after primary infection without clinical symptoms, suggesting bacterial persistence. Yet, information about the induction, regulation and/or location of *C. burnetii* persistence is rare. We have shown that during infection of primary macrophages, hypoxia-induced citrate limitation results in inhibition of *C. burnetii* replication without affecting viability. Here, primary murine macrophages were infected with *C. burnetii* under normoxic (21% O_2_) and hypoxic (0.5% O_2_) conditions to clarify how *C. burnetii* survives this environmental stress condition. Our data suggests that under hypoxic conditions *C. burnetii* does not undergo stringent response, but instead enters a SCV-like form, which is smaller in size and possesses condensed chromatin material and a thicker cell wall. These changes have functional consequences, as the SCV-like persistent form of *C. burnetii* is more infectious, more tolerant to antibiotics and less sensitive to clearance by IFNγ activated macrophages. Hence, the development of the SCV-like persistent form of *C. burnetii* prevents elimination of the pathogen, which in turn allows the pathogen to thrive once the conditions again change in its favor.

## 1. Introduction

*Coxiella burnetii* is a Gram-negative, non-motile, obligate intracellular, pleomorphic coccobacillus [1]. The bacterium is highly resistant to different stresses, including osmotic shock, high temperature, desiccation or UV light, providing environmental stability. This stability is attributed to the small-cell variant (SCV) from, which develops during the biphasic development cycle [2]. The SCV is characterized by a rod-shaped size of 0.2 to 0.5 µm in length (in negative contrast preparations), condensed chromatin and a dense cell wall [3]. Moreover, SCVs display little metabolic activity [2]. Within a host cell the SCV differentiates to the large-cell variant (LCV), which can exceed 1 µm in length (in negative contrast preparations) and represents the replication competent form of *C. burnetii* [3]. Transition from LCV to SCV occurs during the transition from the exponential to the stationary phase of intracellular growth [4].

*C. burnetii* is found worldwide, with the exception of New Zealand. It infects a vast diversity of species ranging from ticks, fish, birds, and rodents to mammals, including ruminants and humans [5]. In humans, it causes the anthropozoonotic disease Q fever [1]. There are two forms of Q fever, acute and chronic Q fever. The acute Q fever mainly represents as flu-like illness, atypical pneumonia or hepatitis. It is either self-limiting or resolves after treatment with doxycycline for two weeks. Chronic Q fever develops months or even years after the initial infection in 2-5% of the cases, suggesting a phase of bacterial persistence prior to the development of disease. The most common manifestation of chronic Q fever represents endocarditis. There are several risk factors for the development of chronic Q fever, such as male gender, preexisting valvular conditions or contact with farm animals. Without treatment chronic Q fever might be fatal. Therefore, early treatment with doxycycline and hydroxychloroquine for at least 18 months is required [6, 7].

To survive adverse conditions, bacteria can enter a state of persistence. Bacterial persistence is transitory in its nature, non-inheritable and quiescent. Importantly, the persistent bacterium returns to normal growth, when the condition changes. Thereby, bacterial persistence is associated with relapsing infections. Bacteria encode different regulatory programs to coordinate persistence, including the general stress response or the stringent response [8-10].

*C. burnetii* DNA was found years after acute Q fever in 65% of the bone marrow aspirates analysed, which might represent the seed for recrudescent infection and the development of chronic Q fever [11]. In addition to bone marrow, adipose tissue was suggested as a site of *C. burnetii* persistence [12, 13]. Neither the location, nor the molecular mechanism(s) resulting in *C. burnetii* persistence are completely understood [9]. Diverse stimuli hinder bacterial replication, such as IFNγ, TNFα or iron limitation [14, 15]. However, whether these stimuli induce *C. burnetii* persistence is unknown. *C. burnetii* possesses genes involved in the general stress response and in the stringent response [16], which might be involved in the development of persistent *C. burnetii*.

Here, we addressed the question whether hypoxia-mediated alteration of the host cell metabolism induces *C. burnetii* persistence. Indeed, our data indicate that under hypoxia, and thus under citrate-limiting conditions, *C. burnetii* establishes an SCV-like persistent form, which is characterized by condensed chromatin, an altered cell wall, increased infectivity and antibiotic tolerance. In addition, we provide evidence that the development of *C. burnetii* persistence is not mediated by the stringent response, but rather by a general stress response.

## 2. Results

### 2.1. Hypoxia triggers a general stress response, but not the stringent response in intracellular C. burnetii

We have previously shown that *C. burnetii* stays viable, but is unable to replicate in hypoxic macrophages [17, 18]. To investigate whether hypoxia-mediated citrate-limitation results in *C. burnetii* persistence, we characterized *C. burnetii* at 48 hours post-infection in normoxic and hypoxic bone marrow-derived macrophages (BMDM) at the transcriptional level. This time point was chosen, as *C. burnetii* has not started to replicate in normoxic BMDM at this time point [17]. Thereby, alterations mediated by replication was omitted. Bacterial persistence is reversible. Consequently, a reoxygenation control (24 hours hypoxia followed by 24 hours normoxia) was included. First, we analysed the expression level of genes involved in the stringent response, which is a ubiquitous stress response to nutrient starvation [19]. A hallmark of the stringent response is the accumulation of the alarmones guanosine tetraphosphate (ppGpp) and pentaphosphate (pppGpp), which are produced by RelA and SpoT [20]. The alarmones together with the transcription factor DksA interact with RNA polymerase to promote binding to the sigma factor RpoS, thereby shifting transcription, which allows the pathogen to adapt to the stress situation [21]. As shown in figure 1a, neither *relA* nor *dksA* are transcriptionally modulated by oxygen limitation. For *spoT* and *rpoS* a reduction of the mRNA level was observed under hypoxic conditions. These data suggest that hypoxia does not induce the stringent response in *C. burnetii*.

**Figure 1.**
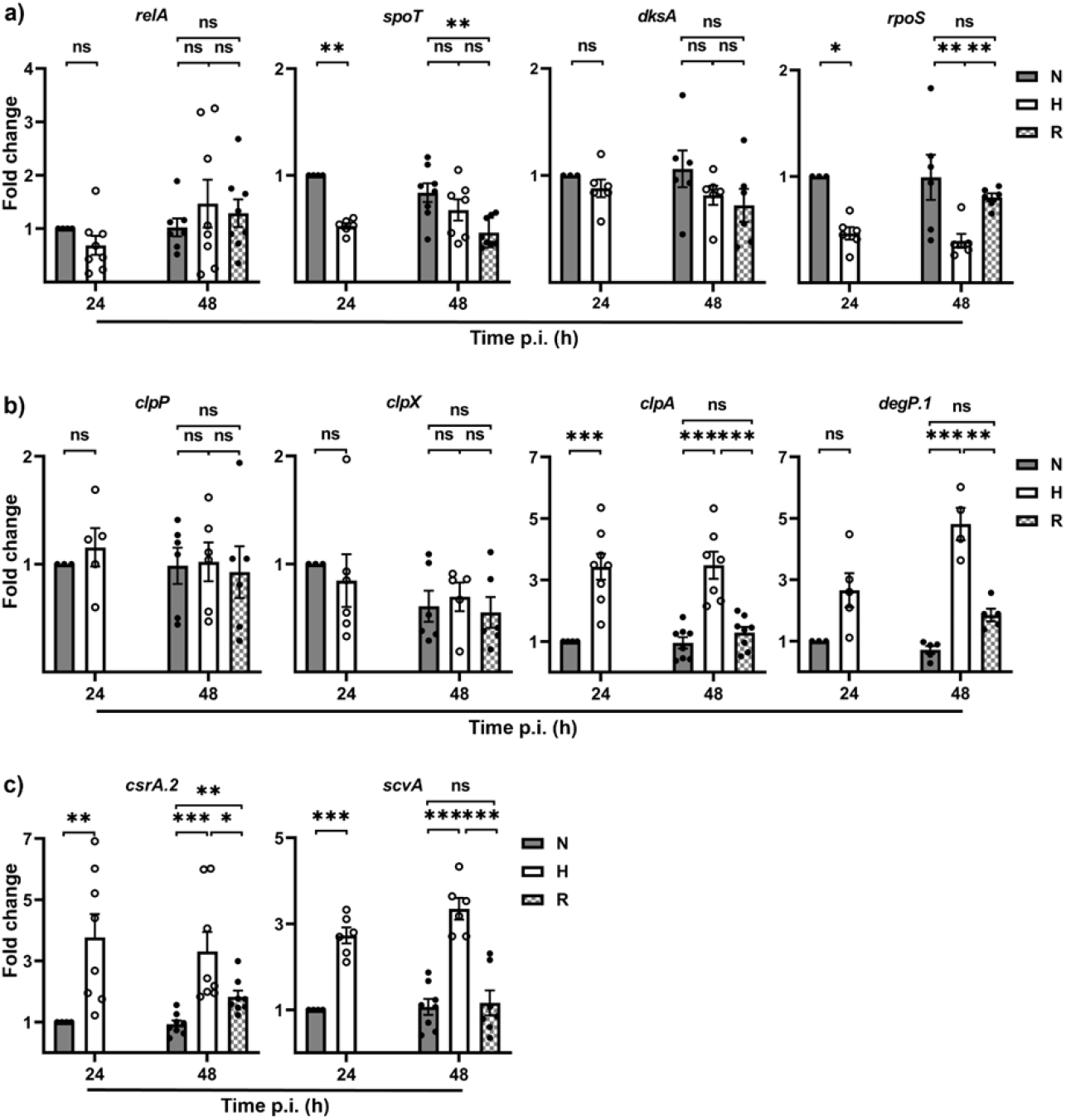
Hypoxia-mediated changes in infected BMDM triggers a general stress response in *C. burnetii*. Murine BMDM were infected for 48 h with *C. burnetii* at a multiplicity of infection of 10 under normoxia (N), hypoxia (H) or re-oxygenated (R) after 24 h of hypoxia. Using qRT-PCR, the gene expression of bacterial genes involved in **a)** the stringent response, **b)** a general stress response and **c)** *scvA* expression was determined. The data is depicted as mean ± SEM of ΔΔCT values (using IS1111 as a calibrator) of 3-4 independent experiments. One-sample t-test and Mann-Whitney U test, ***p<0.001, **p<0.01, *p<0.05, ns=not significant.

As *C. burnetii* lacks a type II toxin-antitoxin system, which is linked to the development of bacterial persistence in other pathogens [9], we concentrated on the expression of genes involved in general stress response. Clp family proteases have multiple functions including regulation of stress responses [22]. *C. burnetii* encodes for the ATPase subunits ClpA and ClpX as well as for the proteolytic subunit ClpP. ClpA and ClpX are chaperones that bind ClpP to establish ClpAP or CLpXP, which allows the efficient processing of large peptides and folded proteins [22]. Similarly, the periplasmic dual-function protease DegP1 is critical for the bacterial stress response by degrading a wide range of proteins [23]. The expression levels of *clpA* and *degP*.*1* were increased in hypoxic BMDM. Importantly, this upregulation was revoked after reoxygenation, demonstrating the reversible nature of this bacterial response (Fig. 1b). The carbon storage regulator (CsrA) binds mRNA and thereby globally regulates gene expression in a posttranscriptional manner [24]. The expression of *csrA*.*2* was upregulated 3 to 4-fold under hypoxic conditions (Fig. 1c). Although the function of CsrA-2 in *C. burnetii* has not been investigated in detail, it has been suggested that it might perform a global regulatory function in the SCV form [25]. Hence, the expression level of *scvA* was determined. ScvA is a small peptide, which is mainly expressed in the SCV form [26]. Indeed, *scvA* is upregulated in infected hypoxic BMDM. After reoxygenation this upregulation was abrogated (Fig. 1c). Together these data indicate that the hypoxic environment within murine BMDM triggers a general stress response rather than a stringent response in *C. burnetii*. Furthermore, the stress induced by the intracellular environment in hypoxic BMDM might trigger the transition from LCV to SCV.

### 2.2. Hypoxia leads to morphological change of intracellular C. burnetii

To determine the developmental stage of *C. burnetii*, the morphology of bacteria in normoxic and hypoxic BMDM was analysed using transmission electron microscopy (TEM). Under hypoxic conditions, BMDM had an increased number of peroxisomes and/or lipid bodies (Fig. 2a), which might indicate a hypoxic stress situation [27]. Both characteristic CCVs (cCCV), that contained predominantly LCVs and only a few SCVs, and abnormal CCVs (aCCV) were seen in normoxic BMDM (Fig. 2a). Under hypoxic conditions, aCCVs were present almost exclusively, often forming multiple smaller CCVs (Fig. 2a). *C. burnetii* in aCCV were often crowded closely together and highly pleomorphic, which did not allow a clear discrimination of LCV and SCV. Thin electron dense bacteria were more frequent, but bacteria with large central rod-shaped electron-dense condensation and swollen bacteria with partly clumped, partly lytic cytoplasm were also seen. The mean diameter of *C. burnetii* in hypoxic BMDM was lower compared to normoxic BMDM indicating a higher frequency of SCV-like forms (Fig. 2b). In addition to the size difference, LCVs and SCVs can be discriminated by the morphology of the bacterial cell wall, especially the periplasmic space (Fig. 2c). As observed in cCCVs under normoxic conditions, the periplasmic space in LCV is electron-lucent and irregularly dilated resulting in an undulating outer membrane, whereas in SCV, it is rather uniform, narrow and filled with an electron dense matrix (Fig. 2d). 36% of the pleomorphic *C. burnetii* in aCCVs under normoxic conditions and 73% of bacteria under hypoxic conditions had SCV-like periplasmic spaces. These data indicate that even normoxic murine BMDM might represent a stressful environment for *C. burnetii*. Importantly, hypoxic conditions boost the development of aCCVs and of the SCV-like form of *C. burnetii*.

**Figure 2.**
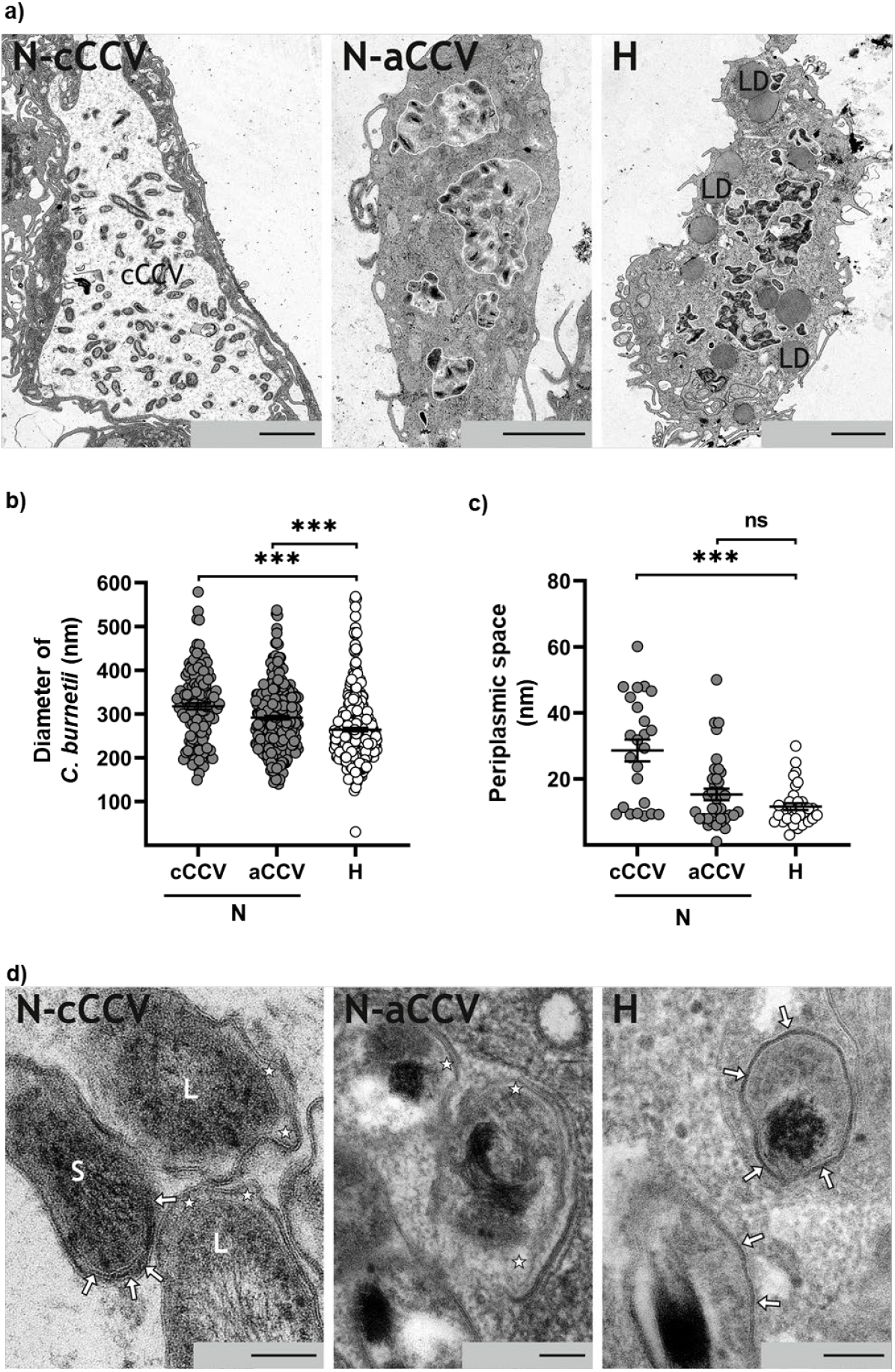
Hypoxia-mediated changes in infected BMDM induce morphological changes in *C. burnetii*. Murine BMDM were infected for 48 h with *C. burnetii* at MOI 10 under normoxia (N) or hypoxia (H). The cells were fixed with ruthenium red glutaraldehyde, embedded in araldite and ultrathin sectioned for electron microscopy. **a)** *C. burnetii* infected normoxic (N) BMDM with a characteristic CCV (cCCV) and another cell with multiple abnormal CCVs (aCCVs), indicated by white outlines and a hypoxic (H) BMDM with multiple aCCVs (indicated by white outlines) are shown. Lipid droplets were marked with LD. The scale bar = 2 µm. **b)** The diameter of 140 bacteria in cCCV and 300 bacteria in aCCV of normoxic and 300 bacteria in hypoxic BMDM were measured. The data is depicted as mean ± SEM from 2-3 independent experiments. Mann-Whitney U test, ***p<0.001. **c)** The maximal thickness of the periplasmic space was measured from at least 30 bacteria of each condition. Shown are the means ± SEM, Mann-Whitney U test, ***p<0.001. **d)** Morphology of bacterial wall of *C. burnetii* in BMDM cells is shown: two LCVs (L) with irregularly dilated electron lucent periplasmic spaces (*) and one SCV (S) with narrow, electron dense periplasmic space (arrows) in a cCCV under normoxic conditions (N-cCCV), three pleomorphic *C. burnetii* with LCV-like periplasmic spaces (*) in an aCCV under normoxic conditions (N-aCCV), and two bacteria with SCV-like periplasmic spaces (arrows) under hypoxic conditions. The scale bar = 100 nm.

To verify that the development of SCV-like *C. burnetii* is not only restricted to hypoxic murine BMDM, *C. burnetii* grown in the human endothelial cell line EA.hy926 were analysed. As in BMDM, increased numbers of peroxisomes and occasionally lipid bodies indicated hypoxic stress situation (Fig. 3a). Normoxic EA.hy926 cells contained large CCVs with few to moderate numbers of bacteria, predominantly LCVs and rarely SCVs. In hypoxic EA.hy926 cells, the morphologies of the CCVs and of *C. burnetii* differed markedly. Bacteria were distributed in multiple small aCCVs throughout the cells as described in BMDM (Fig. 3a). A membrane limiting aCCVs was not always clearly detected (Fig. 3a). The bacteria within aCCV had highly variable morphology, but the size was comparable to SCVs. In detail, *C. burnetii* in normoxic cells had a mean diameter of 376 nm ± 34 nm, while the diameter of bacteria in hypoxic cells was 210 nm ± 21 nm (Fig. 3b). In addition, a difference in the size and morphology of the periplasmic space was obvious between cells under normoxic and hypoxic conditions (Fig. 3c and d). The majority of bacteria in normoxic endothelial cells were LCVs and, thus, had the characteristic electron lucent and irregularly dilated periplasmic space. Irrespective of their heterogenous morphology, the periplasmic spaces of *C. burnetii* in hypoxic cells were comparable to those in SCVs (Fig. 3d). The data indicate that the changes induced by hypoxia are more clearly seen in EA.hy926 cells (Fig. 3) compared to BMDM (Fig. 2). Nevertheless, the data demonstrate that intracellular *C. burnetii* enter an SCV-like phenotype under hypoxic conditions, independently of the host species or cell type.

**Figure 3.**
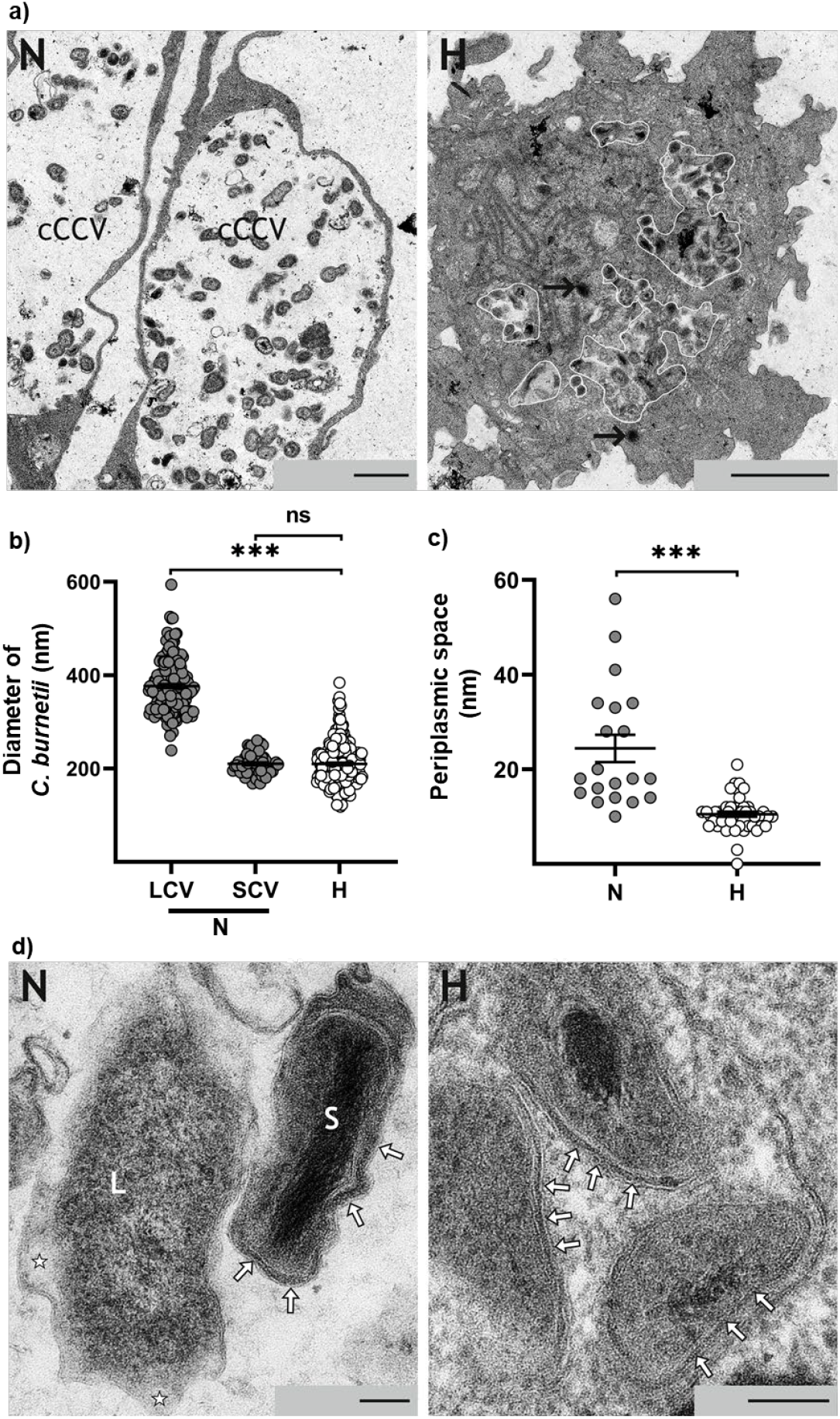
Hypoxia-mediated changes in infected human endothelial cells induce morphological changes in *C. burnetii*. Human EA.hy926 cells were infected for 48 h with *C. burnetii* at MOI 200 under normoxia (N) or hypoxia (H). The cells were fixed with ruthenium red glutaraldehyde, embedded in araldite and ultrathin sectioned for electron microscopy (EM). **a)** Two *C. burnetii* infected normoxic (N) cells with large cCCV and a hypoxic (H) EA.hy926 cell with multiple small aCCV (indicated by white outlines) are shown. Peroxisomes were highlighted by arrows. The scale bar = 2 µm. **b)** The size of 250 bacteria per conditions from 2-3 independent experiments were measured. The data is depicted as mean ± SEM. Mann-Whitney U test, ***p<0.001. **c)** The thickness of the periplasmic space was measured from 20 bacteria (10 LCV and 10 SCV) under normoxic and 40 bacteria under hypoxic conditions. Shown are the means ± SEM, t-test. ***p<0.001. **d)** Morphology of the bacterial wall of *C. burnetii* EA.hy926 cell is shown: one LCV (L) with irregularly dilated electron lucent periplasmic space (*) and one SCV (S) with narrow, electron dense periplasmic space (arrows) under normoxic (N) conditions and three pleomorphic bacteria with SCV-like periplasmic spaces (arrows) under hypoxic conditions. The scale bar = 10 nm.

### 2.3. Morphological changes of SCV-like persistent C. burnetii were accompanied by upregulation of cell wall remodelling enzymes

The expression of genes encoding proteins involved in cell wall remodelling was shown to be upregulated during the late stationary phase, at a time point of transition to the SCV form [28]. Hence, the expression of such genes was analysed to determine the underlying reason for the morphological changes in the cell wall of SCV-like persistent *C. burnetii* in hypoxic BMDM. The expression of the YukD domain protein containing gene *enhA*.*2*, the lytic murein transglycosylase B encoding gene *lmtg-ß* and *enhC* were upregulated under hypoxic conditions (Fig. 4). Reoxygenation reset the hypoxia-induced upregulation of *enhA*.*2, lmtg-ß* and *enhC*, demonstrating the transient nature of this transcriptional regulation (Fig. 4).

**Figure 4.**
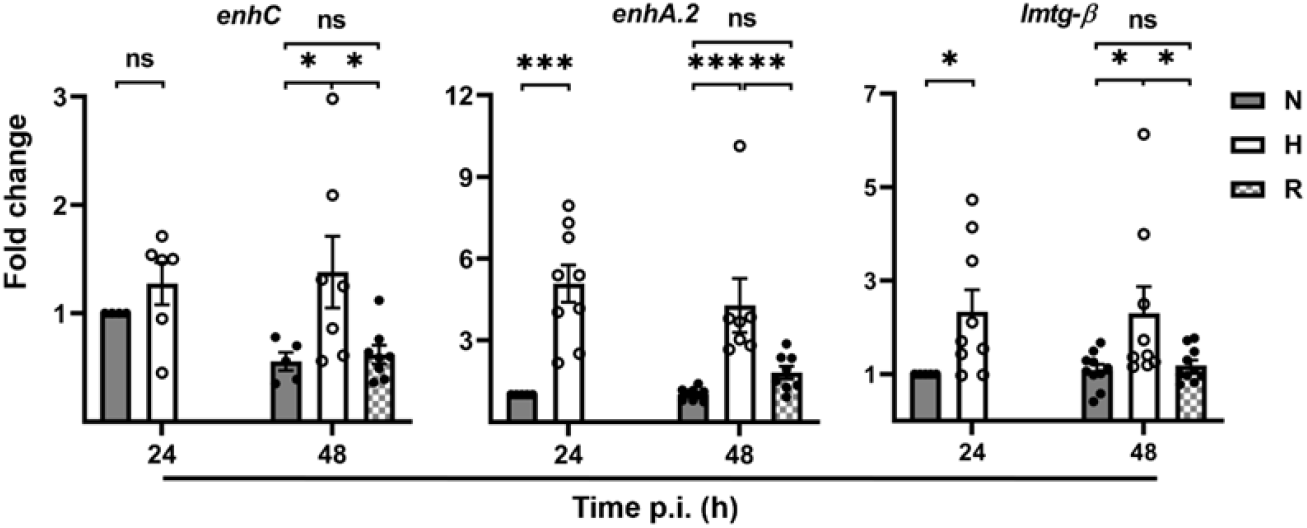
Hypoxia-mediated changes in infected BMDM induce upregulation of *C. burnetii* genes encoding cell wall remodelling proteins. Murine BMDM were infected for 48 h with *C. burnetii* at MOI of 10 under normoxia (N) or hypoxia (H). Using qRT-PCR, the gene expression of bacterial proteins involved in cell wall remodelling was determined. The data is depicted as mean ± SEM of □□CT values (using IS1111 as a calibrator) from 3-4 independent experiments. One-sample t-test and Mann-Whitney U test, ***p<0.001, **p<0.01, *p<0.05, ns=not significant.

### 2.4. The SCV-like persistent C. burnetii are more infectious and resistant to IFNg treatment

The SCV is often reported as the infectious form of *C. burnetii* [28]. Consequently, the infectivity of bacteria isolated from normoxic or hypoxic BMDM were determined. CHO cells were infected for 4 hours with *C. burnetii* isolated from normoxic or hypoxic BMDM and the infection rates were determined by CFU assay. The numbers of intracellular bacteria were higher when CHO cells were infected with *C. burnetii* isolated from hypoxic BMDM (Fig. 5a). This result indicates that SCV-like persistent *C. burnetii* are more infectious.

**Figure 5.**
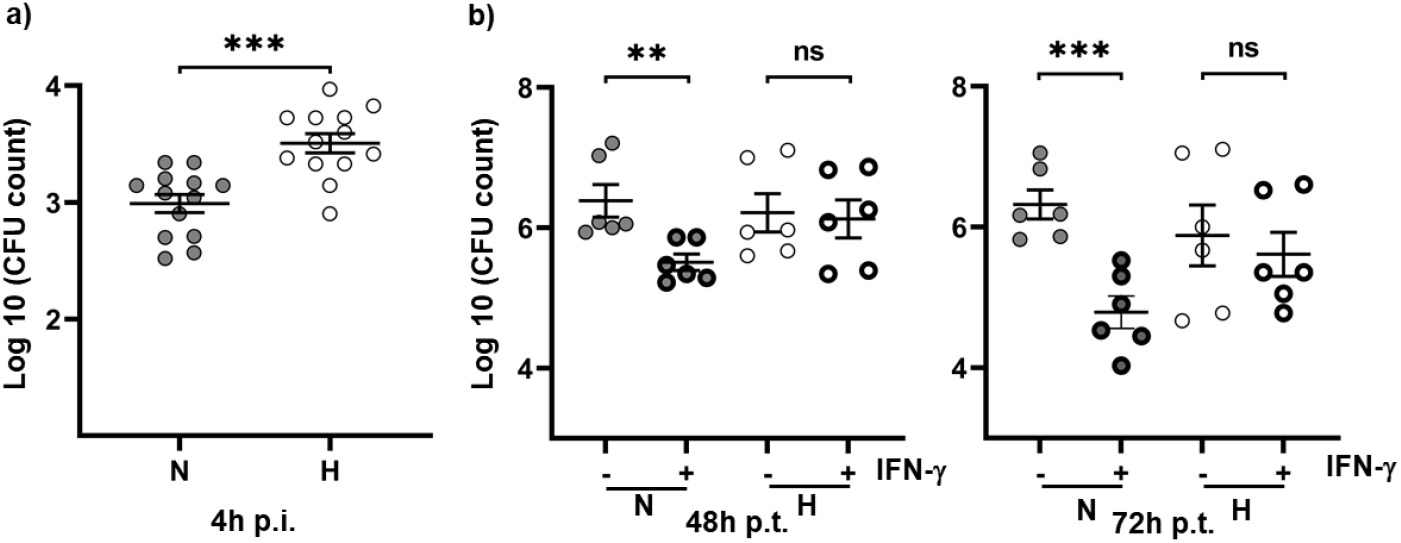
SCV-like persistent *C. burnetii* are characterized by increased infectivity. Murine BMDM were infected for 48 hours with *C. burnetii* at a MOI of 10 under normoxia (N) or hypoxia (H). **a)** *C. burnetii* were isolated and used to infect CHO cells for 4 h. The infection rate was determined by colony forming unit (CFU) counts from 3-4 independent experiment. The data depicted as means ± SEM. T-test, ***p<0.0001. **b)** Infected cells were treated with 10 ng/ml IFNγ for 48 and 72 h under normoxic conditions. CFU counts were performed and survival was determined. Results from three independent experiments are shown. Mann-Whitney U and t-test, ***p<0.001, **p<0.01, ns=not significant.

Cell autonomous control is important to control and clear the infection. It was shown that IFNg-activation results in efficient cell autonomous control of a *C. burnetii* infection [29, 30]. Therefore, the consequence of IFNγ activation on the survival of *C. burnetii* in normoxic and hypoxic BMDM was analysed. Importantly, the infected normoxic and hypoxic BMDM were transferred to normoxic conditions for IFNg treatment. While treatment with IFNγ resulted in killing of intracellular *C. burnetii* by previously normoxic BMDM over time, this was not observed in BMDM kept previously under hypoxic conditions (Fig. 5b). These data suggest that the SCV-like persistent bacteria are less sensitive to IFNγ-activated cell autonomous control.

### 2.5. The SCV-like persistent C. burnetii are characterized by increased tolerance to carbenicillin and doxycycline

Persistent bacteria become refractory to antibiotic treatment [31]. Therefore, the antibiotic susceptibility of *C. burnetii* was investigated. *C. burnetii* isolated from normoxic or hypoxic BMDM were treated in axenic culture with either doxycycline or carbenicillin [32]. Doxycycline was chosen as it is the first-line treatment of acute Q fever. Carbenicillin was used as it has been shown that a thinner cell wall/outer membrane complex resulted in increased susceptibility to this antibiotic [16], suggesting that SCV-like persistent *C. burnetii* might be more tolerant to this carboxypenicillin. Indeed, *C. burnetii*, isolated from hypoxic BMDM, are more resistant to treatment with carbenicillin (Fig. 6a), but also slightly, but significantly, to doxycycline (Fig. 6b). These data indicate that the morphological changes observed in the SCV-like persistent *C. burnetii*, resulted in reduced susceptibility to the antibiotics doxycycline and carbenicillin.

**Figure 6.**
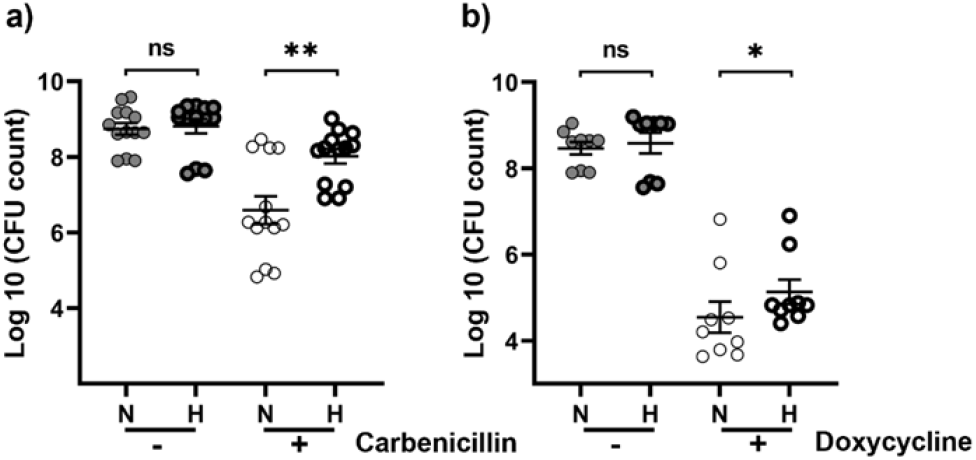
SCV-like persistent *C. burnetii* are more antibiotic tolerant. Murine BMDM were infected with *C. burnetii* at a MOI of 10 under normoxia (N) or hypoxia (H). 48 h post-infection *C. burnetii* were isolated and treated with **a)** 150 µg/ml carbenicillin or **b)** 2ng/ml doxycycline in ACCM-2 medium for 7 days at 2.5% O_2_, 5% CO_2_ and 37°C. CFU was performed from 3-5 independent experiments. Mann-Whitney U test, **p<0.01, *p<0.05, ns=not significant.

## 3. Discussion

Authors should discuss the results and how they can be interpreted from the perspective of previous studies and of the working hypotheses. The findings and their implications should be discussed in the broadest context possible. Future research directions may also be highlighted. Bacterial persistence is the cause for reoccurring and relapsing infections, as persistent bacteria are not sufficiently cleared by the host immune system [8]. This can be due to a lack of recognition by the immune system, mediated by e.g. cell wall modulation [33]. In addition, persistent bacteria are often insensitive towards antibiotics, which prevents successful treatment [34]. This allows survival of the bacteria, which in turn allows their expansion, once the cause for bacterial persistence is resolved, and concomitant disease progression.

Our knowledge about *C. burnetii* persistence is limited, although the long time-span between infection and disease outbreak indicates that *C. burnetii* persistence might be an important step towards chronic Q fever. A few reports have suggested that *C. burnetii* may persist in bone marrow [11] or in adipose tissue [12]. Importantly, at both sites the oxygen level might be reduced [35, 36]. This and the fact that *C. burnetii* survives, but does not replicate in hypoxic macrophages [17], promoted us to investigate the physiological state of these bacteria in more detail. Of note, the low level of oxygen did not prevent *C. burnetii* replication in hypoxic BMDM *per se*, but rather the hypoxia-induced modulation of host cell signalling and metabolism [17]. Nutrient limitation triggers the stringent response in bacteria to reprogram from rapid growth to metabolic homeostasis [8]. Although we observed a lack of the TCA cycle metabolite citrate as being responsible for the growth defect of *C. burnetii* in hypoxic BMDM [17], we did not observe transcriptional upregulation of the stringent response under hypoxic conditions (Fig. 1a). Instead, we observed a general stress response (Fig. 1b) and increased expression of *scvA* (Fig. 1c). While the function of ScvA in *C. burnetii* is not completely known, it is an important marker for the development of SCVs. The expression of *scvA* is directly regulated by RpoS. Thus, *scvA* transcription is downregulated in D*rpoS C. burnetii* [16]. However, under hypoxic conditions *rpoS* is downregulated, while *scvA* is upregulated (Fig. 1a and c), suggesting that other factors also regulate its expression. Noteworthy, *C. burnetii* enters a SCV-like form under hypoxic conditions (Figs. 1c, 2 and 3). So far, the SCV has been described as the extracellular stable form required for environmental transmission [2, 26]. The intracellular transition to the SCV occurs in the stationary phase approximately six days post-infection [37], a time point correlating with bacterial egress [38]. Our data suggest that the SCV is not only important for the extracellular phase of the pathogen, but also to survive adverse intracellular conditions. Interestingly, the characteristics of the CCVs containing the SCV-like persistent *C. burnetii* differs quite significant from the classical described CCV. The aCCVs were more pleomorphic and smaller in size. The fact that multiple aCCVs were present in one cell (Fig. 2a and 3a) indicates that aCCVs did not undergo homotypic fusion. While the underlying reason is unknown, it can be speculated, that the SCV-like persistent bacteria did not assemble the T4BSS [39], which translocates effector proteins involved in homotypic fusion events [40, 41].

The morphological changes observed (Figs. 2 and 3) seem to allow the pathogen to survive intracellularly. How these changes are precisely regulated is unknown. It has been speculated that ScvA plays a role in chromatin condensation, which protects DNA from stress-induced insults [26]. Consequently, the DNA condensation observed in the SCV-like persistent *C. burnetii* (Figs. 2d and 3d) might be due to upregulation of *scvA* (Fig. 1c). Similarly, the alterations in the cell wall morphology observed in SCV-like persistent *C. burnetii* (Figs. 2c, 2d, 3c and 3d) might be caused by increased expression of *enhC, enhA*.*2* and/or *lmtg-ß* (Fig. 4). Further research will be required to disentangle the signalling cascade(s) leading to modulation of bacterial morphology. These changes might cause rigidity of the cell wall, allowing *C. burnetii* to survive either host cell autonomous defence (Fig. 5b) or antibiotic treatment (Fig. 6a and b). Specifically, the modulation of the cell wall structure might be relevant for the latter. Peptidoglycan, the major component of the bacterial cell wall, is a net-like heteropolymer made of glycan chains crosslinked by short peptides [42, 43]. Transpeptidases, including penicillin-binding proteins (PBPs), are enzymes which crosslink the peptidoglycan chains. The majority of peptide crosslinks are of the 4,3 type and are catalysed by DD-transpeptidases [43]. However, bacteria also encode for LD-transpeptidases, which present a YkuD-like domain, and produce a 3,3 crosslink [44]. Both crosslink types serve to strengthen the cell wall. In *Escherichia coli* the LD-transpeptidases are upregulated in the stationary phase and under stress conditions [45]. EnhA.2 is a YkuD-like domain-containing protein [28] and is upregulated under intracellular hypoxic conditions (Fig. 4). Importantly, the SCV contains higher percentage of 3,3 crosslinks [28], indicating that upregulation of cell wall remodelling enzymes (Fig. 4), including EnhA.2, might result in increased 3,3 crosslinking. Bacteria containing 3,3 crosslinks were shown to be more resistant to beta-lactams [46] and to have improved cell envelope integrity [45]. Thus, the increased 3,3 crosslinks might result in improved stability of the cell wall, which in turn might provide improved antibiotic tolerance (Fig. 6a and b). Further investigations are necessary to prove this assumption. This might be relevant as the switch from 4,3 to 3,3 crosslinking might be a new therapeutic target to improve antibiotic susceptibility of this obligate intracellular pathogen.

## 4. Materials and Methods

### Reagents

Chemicals were purchased from Sigma Aldrich unless indicated otherwise.

### Mice

C57BL/6 wild-type mice were obtained from Charles River Laboratories.

### Isolation and Differentiation of Murine Bone Marrow Derived Macrophages

Femurs and tibiae were isolated from C57BL/6 wild-type mice. The bone marrow cavity was flushed with cMoAb medium containing RPMI 1640 (Thermo Fisher), 10% fetal calf serum (FCS), 1% HEPES (AppliChem), 0.5% ß-mercaptoethanol using a 10 ml syringe fitted with a 27G needle. The cells were centrifugated at 365g for 10 min. The pellet was resuspended in macrophage differentiation medium (MDM) composed of DMEM-GlutaMAX (Gibco), 15% FCS, 20% supernatant from L929 cells, 1% penicillin/streptomycin (P/S, Gibco), 1% Non-essential amino acids (NEAA, Gibco) and 0.5% HEPES. 7×10^6^ cells were seeded per T75 culture flasks (Thermo Fisher) and differentiated for 7 days at 37°C and 5% CO2. The culture was supplemented with an additional 10 ml MDM on day 4.

### Cells and cell culture

To harvest the differentiated BMDMs, the culture medium was removed and ice-cold PBS was added. Cell scrapers were used to detach the cells from the culture bottle. The cell suspension was centrifuged at 365g for 10 min and the pellet resuspended in cMoAb medium at a concentration of 5×10^5^ cells/ml.

The human endothelial cell line EA.hy926 (ATCC®CRL-2922^™^) was cultivated in DMEM-GlutaMAX and 5% FCS. The CHO-FcR cells were cultivated in DMEM/F-12 (Thermo Fisher) and 5% FCS.

### Culture of *Coxiella burnetii*

*Coxiella burnetii* NMII, clone 4 were inoculated at a concentration of 10^6^/ml in ACCM-2 (Sunrise Science Products) and cultivated for five days at 37°C, 5% CO2 and 2.5% O2. The bacterial culture was centrifuged at 4309g for 20 min and the pellet resuspended in PBS. The bacterial concentration was determined spectrophotometrically at OD600, with an OD of 1 corresponding to 10^9^ bacteria per ml. For infection experiments, a multiplicity of infection (MOI) of 10 was used for BMDMs, while a MOI of 200 was used for EA.hy926 cells.

### Hypoxia

For maintaining a hypoxic environment, experiments were performed within an InvivO2 hypoxic chamber (Baker Ruskinn) to control oxygen levels. Cells were treated under a defined hypoxic environment of 0.5% O2, 5% CO2, and 37 °C. To ensure complete deoxygenation, all buffers and media were pre-incubated in the hypoxic chamber for a minimum of 4 h before being applied to the hypoxic cells.

### Infection of BMDM

The BMDMs were infected with *C. burnetii* at a MOI of 10 for 4 h under normoxic (21% O2, 5% CO2 and 37 °C) and hypoxic (0.5% O2, 5% CO2 and 37 °C) conditions. Afterwards, the cells were washed thrice with 1x PBS to eliminate extracellular bacteria. The cells were then cultured for additional 48 h in fresh pre-incubated cMoAb medium.

### Infection of EA.hy926 cells

5×10^4^ EA.hy926 cells/ml were seeded 18 hr prior to infection with *C. burnetii* at MOI of 200. The infected cells were incubated for 4 h under normoxic (21% O2, 5% CO2 and 37 °C) or hypoxic conditions (0.5% O2, 5% CO2 and 37 °C). Afterwards, the cells were washed thrice with 1x PBS to eliminate extracellular bacteria. The EA.hy926 cells were subsequently cultured for an additional 24 h or 48 h in fresh, pre-incubated DMEM-GlutaMAX medium supplemented with 5% FCS.

### RNA extraction, cDNA synthesis and qRT-PCR

The cells were lysed using 500 µl TRIzol reagent. RNA isolation was performed using the Direct-zol™ RNA Mini Prep Plus kit (Zymo research) according to the manufacturer’s instructions. To remove contamination of genomic DNA, a DNase treatment was performed using the RapidOut DNA removal kit (Thermo Fisher Scientific), according to manufacturer’s instructions. cDNA was synthesized from 500 ng to 1 µg purified RNA by using the High-Capacity cDNA Reverse Transcription Kit (Applied Biosystems), according to the manufacturer’s instructions. Quantitative real-time PCR (qRT-PCR) was carried out using the PowerUp™ SYBER™ Green Master Mix (Applied Biosystems), following the manufacturer’s instructions. The cDNA was diluted 1:5 and the primers (100 nM final concentration) listed in Table 1 were used. The quantification of the expression levels of the genes was referenced to the housekeeping gene IS1111 and normalized to the normoxic sample. The 2^-ΔΔCT^ method was used to calculate the fold change.

**Table 1.**
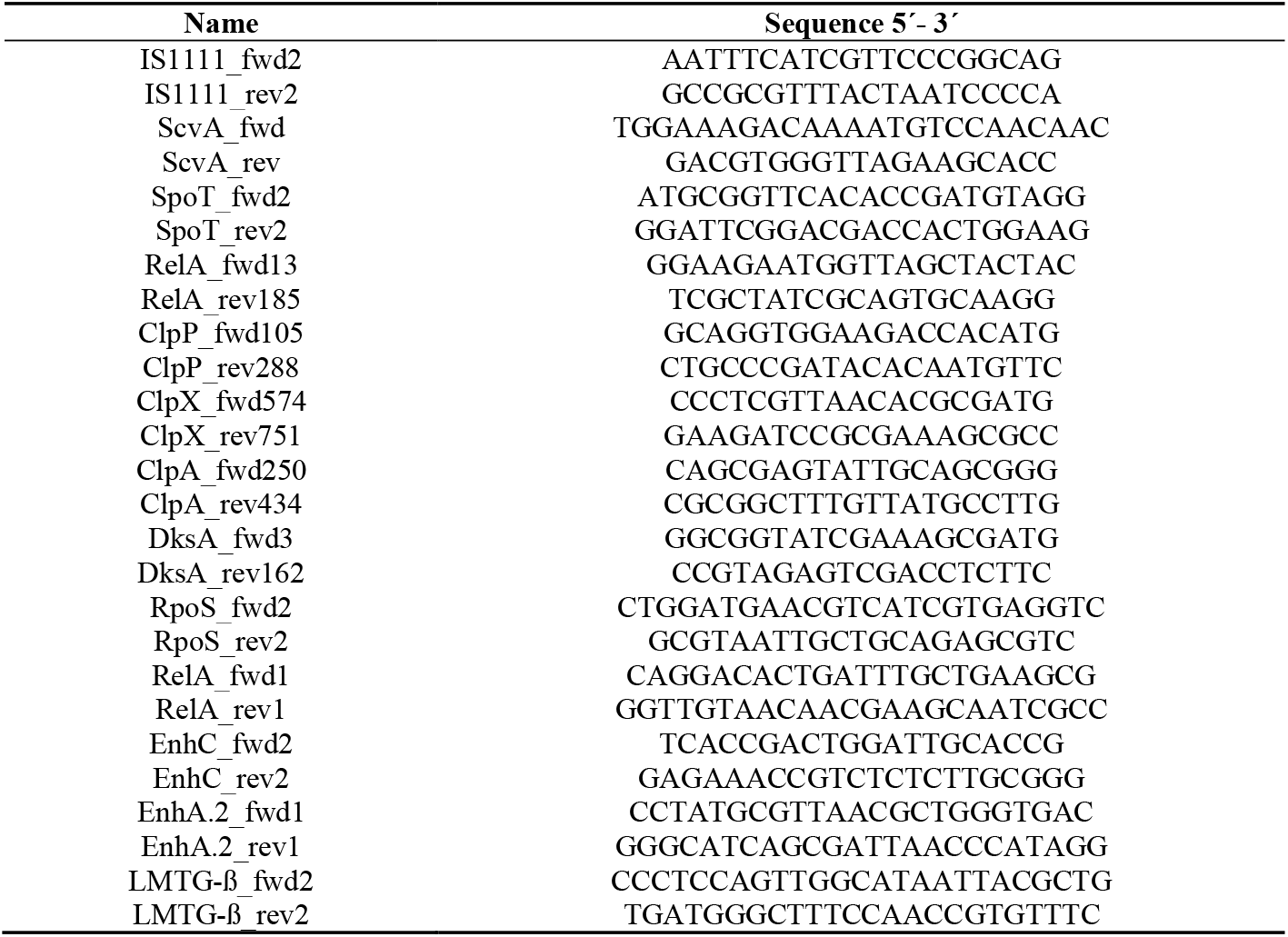
Primers used.

### Colony forming unit assay (CFU)

The medium was removed and the infected cells washed washing thrice with 1x PBS. The cells were shocked with 1 ml of ice-cold ddH2O and kept at 4 °C for 30 min. Cell lysis was executed by vigorous pipetting. The bacteria were pelleted by centrifugation at 20,817g for 1 min and the pellet was resuspended in 200 µl PBS. A 10-fold serial dilution was carried out until a dilution factor of 10^-5^ was reached. From each dilution, 10 µl were spotted in triplicates onto ACCM-2/agarose plates containing 0.3% agarose. The plates were incubated for 8-10 days at 2.5% O_2_, 5% CO_2_ at 37 °C. Colonies were counted microscopically.

### IFNγ treatment

BMDMs were infected with *C. burnetii* for 48 h under normoxic and hypoxic conditions. The medium was replaced with fresh medium with or without 10 ng/ml IFNγ. The cells were incubated under normoxic conditions for 48 h and 72 h. An CFU assay was performed to determine the numbers of viable bacteria.

### Infectivity assay

Bacteria were isolated 48 h post-infection from normoxic and hypoxic BMDM by osmotic lysis. In detail, the infected cells were washed thrice with PBS, shocked with 1 ml of ice-cold ddH2O for 10 min at RT and and 30 min at 4 °C. Cell lysis was executed by vigorous pipetting. The bacteria were pelleted by centrifugation at 20,817g for 1 min and the pellet was resuspended in 200 µl PBS. 10 µl of isolated bacteria from each condition were used to infect CHO cells for 4 h. An CFU assay was performed to evaluate the infection rate.

### Antibiotic treatment

Isolated bacteria were resuspended in 300 µl of 1x ACCM-2. 100 µl each were added to 4 ml 1x ACCM-2 medium with and without antibiotic (3 ng/ml doxycycline or 150 µg/ml carbenicillin). After 7 days of incubation at 2.5% O_2_, 5% CO_2_ at 37 °C, a CFU assay was performed.

### Electron microscopy sample preparation (EM)

For electron microscopy, 1×10^6^ BMDMs and 2×10^5^ EA.hy926 cells were seeded in 6 well plates. The cells were infected at a MOI of 10 (BMDMs) and 200 (EA.hy926) for 48 h under both hypoxic and normoxic conditions. 48 h post infection, the cells were fixed by adding 5 ml of ice-cold ruthenium red-glutaraldehyde fixative in 0.1M cacodylate buffer (pH 7.3, 2.5% glutaraldehyde (Roth, Karlsruhe), 2% paraformaldehyde (Roth, Karlsruhe), 0.075% ruthenium red (Serva, Heidelberg), cacodylate acid (Roth, Karlsruhe)) per well and incubated for 3 h at 4 °C. Gentle agitation was applied intermittently during fixation to ensure uniform exposure. The fixative was removed and the cells washed three times for 15 min with ice-cold 0.1 M cacodylate buffer. The cells were carefully scraped from the wells and centrifuged at 1500g for 5 min at 4 °C. The cell pellets were resuspended and stored in cacodylate buffer at 4 °C until further processing.

Samples were centrifuged at 1500g for 5 min, the supernatant discarded and the resulting cell pellets mixed with a drop of liquid 2% agarose (50 °C) (Roth, Karlsruhe). After cooling, the agarose containing the cells was cut in 2 mm^3^ blocks and transferred to cacodylate buffer (pH 7.3). The agarose blocks were embedded in araldite CY212 (Plano, Wetzlar) using a tissue processer (Lynx II, Electron Microscopic Sciences, Hatfield, USA). A 2 h treatment with osmic acid-ruthenium red solution (2% osmic acid in aqua bidest (Roth, Karlsruhe), 0.06% ruthenium red in 0.1M cacodylate buffer) was included before dehydration for post-fixation of lipid membranes. Acetone (Roth, Karlsruhe) was used for dehydration. The incubation steps were performed under vacuum and with mild agitation over 28 h. At the end, the agarose blocks were transferred to tissue moulds with freshly prepared araldite CY212 and cured at 50 °C for 48 h and at 65 °C for 24 h. Excessive resin was trimmed off (Leica-EM-Trim, Leica, Wetzlar). Semi-thin sections of 0.3 µm thickness were cut on a ultramicrotome (Leica Ultracut UCT, Leica, Wetzlar) using a diamond knife (Histo, Diatome, Nidau, Switzerland) and stained with 0.25 % toluidine blue. They were used to select defined areas for ultrathin sectioning. Ultra-thin sections of 80 nm thickness were cut on a ultramicrotome using a diamond knife (Ultra 35°, Diatome, Nidau, Switzerland), collected on 300-mesh copper grids (Plano, Wetzlar) and contrasted with lead citrate (Leica, Wetzlar) and uranyl acetate (Plano, Wetzlar) using an automatic stainer (Leica EM AC20, Leica, Wetzlar). Samples were examined in a transmission electron microscope (TECNAI 12, FEI Deutschland, Dreieich) at an acceleration voltage of 80 KV, and findings documented using a digital camera (TEMCAM FX416, TVIPS, Gauting). Measurements were performed on the ultra-micrographs using the EM-Measure software (version from 2021-05-19, TVIPS, Gauting). Micrographs were optimized for brightness and contrast using Adobe Lightroom® (Adobe CS, Version 2025). Image sections were selected, scale bars set using Adobe Indesign® (Adobe CS, Version 2025). Final images were served as sRGB-tif files using Photoshop® (Adobe CS, Version 2025).

### Statistical analysis

Statistical analyses were conducted using GraphPad Prism (version 10). Statistical details can be found in the legends of the figures. Normality distribution was tested with D’Agostino & Pearson normality test or the Kolmogorov-Smirnov test, or, when N < 8, Shapiro-Wilk normality test or the Kolmogorov-Smirnov test. For non-normally distributed datasets, the Mann-Whitney U test was used. Otherwise, the unpaired two-tailed Student’s t test was used. One-sample t-test was used when comparing datasets to normalized values. A value of p < 0.05 was considered significant.

## Author Contributions

Conceptualization, A.L. and I.H.; methodology, E.L.T.; formal analysis, F.A. and I.H..; investigation, F.A. and I.H.; resources, A.L., C.B. and E.L.T.; data curation, F.A., I.H. and E.L.T..; writing—A.L.; writing—review and editing, F.A., I.H., C.B., and E.L.T.; visualization, F.A. and E.L.T.; supervision, A.L.; project administration, A.L.; funding acquisition, A.L. All authors have read and agreed to the published version of the manuscript.

## Acknowledgments

We thank Kevin Hause for preparation of the EM samples and Marcus Pfau for photographic work and editing of the EM images.

